# Stress Granule Coarsening Is a Pathological Inflection Point for Cardiac Electrophysiological Dysfunction

**DOI:** 10.64898/2026.06.19.733463

**Authors:** Heather L. Struckman, Isabelle Field, Anna Z. Li, Erica Marquez, Martina M. Seidel, Thomas M. Schuster, Caroline Chou, Samantha Giangrasso, Kory J. Lavine, Dzmitry Matsiukevich, David M. Ornitz, Nathaniel Huebsch, Anastasia Khokhlova, Jonathan R. Silva

**Affiliations:** Department of Biomedical Engineering, McKelvey School of Engineering, Washington University in St. Louis, St. Louis, MO, US; Department of Medicine, Division of Cardiology, Washington University in St. Louis School of Medicine, St. Louis, MO, US; Department of Developmental Biology, Washington University in St. Louis School of Medicine, St. Louis, MO, US; Department of Pathology and Immunology, Washington University in St. Louis School of Medicine, St. Louis, MO, US; Department of Pediatrics, Washington University in St. Louis School of Medicine, St. Louis, MO, US

## Abstract

Arrhythmia risk rises early in many forms of cardiac stress, often before contractile failure is evident. Stressed cardiomyocytes accumulate biomolecular condensates, known as stress granules (SGs), whose contribution to this electrical vulnerability has been unclear. We mapped where SGs reside and followed their life cycle under acute and chronic oxidative stress across complementary model systems and assessed electrophysiological consequences with pharmacological tools targeting granule assembly, microtubule integrity, and calcium channel function. Under both stress regimes, SGs localized preferentially to z-lines and intercalated discs, marking these mechanically critical sites as hubs of condensate assembly. Merging of granules, referred to as coarsening, rather than initial formation, emerged as a pathological connection. Early granules were broadly cytoprotective, whereas progressive coarsening was accompanied by disruption of alpha-actinin and L-type calcium channel (Cav1.2) nanodomains and by shortening of action potential (AP) duration. Coarsened granules disorganized Cav1.2 nanodomains through a microtubule-dependent mechanism, and arresting coarsening with nocodazole preserved nanodomain integrity and restored AP morphology. The transition from nascent to coarsened SGs therefore represents a targetable inflection point, and limiting coarsening may prevent proarrhythmic remodeling during cardiac oxidative stress.

## Introduction

Stress granules (SGs) are membrane-less, cytoplasmic biomolecular condensates that form in response to cellular stress. They are defined as the cytoplasmic foci at which untranslated mRNAs accumulate.^1,2^ SGs arise from pools of untranslated mRNA when cellular homeostasis is disrupted, forming cytoplasmic mRNP granules.^3–5^ Their formation proceeds through two consecutive steps. First, cellular stress activates one of four integrated stress response kinases (protein kinase R (PKR), protein kinase RNA-like endoplasmic reticulum kinase (PERK), general control nonderepressible 2 (GCN2), or heme-regulated inhibitor (HRI)), which phosphorylates the alpha subunit of eukaryotic initiation factor 2 (eIF2). This inhibits translation initiation and triggers polysome disassembly.^6^ Second, the resulting stalled mRNP complexes, together with scaffold proteins such as G3BP stress granule assembly factor 1 (G3BP1), G3BP stress granule assembly factor 2 (G3BP2), and T-cell intracellular antigen-1 (TIA-1), coalesce into discrete cytoplasmic foci through liquid-liquid phase separation. As local protein concentrations exceed the phase separation threshold, these assemblies stabilize into mature granules.^6–9^

Initial SG formation is protective, as granules limit cellular damage by sequestering non-translating mRNA pools and redirecting resources toward stress resolution and survival. Acute SG assembly is broadly cytoprotective, whereas chronic SG persistence becomes pathological.^10–12^ The transition between these states is governed by granule size. Newly formed small granules are transported along microtubules by dynein and kinesin motors and progressively fuse into larger assemblies through a process termed coarsening. While microtubules are dispensable for initial SG nucleation they are required for the secondary coalescence step; their disruption arrests granule growth without affecting nucleation.^13–16^ Dynein, which drives cargo transport toward the microtubule minus-end, is essential for SG assembly and coarsening, as its inhibition leads to smaller and fewer granules.^17–19^ In contrast, kinesin participates in SG disassembly and limits granule growth and its depletion results in persistent granule accumulation.^14,20^ When stress resolves, small granules are cleared through chaperone-mediated pathways; however, sustained stress can overwhelm these systems, causing disassembly to fail and driving a liquid-to-solid transition that underlies pathological granule behavior.^21,22^ Large persistent granules that escape chaperone resolution require autophagy and proteasome-dependent degradation, and their accumulation is associated with the formation of aberrant disease-linked structures.^9,21,23^ Thus, granule size and clearance dynamics, rather than formation alone, determine whether SGs function as protective translational hubs or drivers of pathological remodeling.

SGs have been reported across a range of cardiac pathologies, implicating their assembly factors in both chronic and acute disease. In chronic cardiac remodeling, G3BP1 expression is upregulated during hypertrophy, where it promotes hypertrophic translation and regulates the hypertrophic transcriptome. Knockdown of G3BP1 restricts cardiomyocyte growth and suppresses gene programs associated with contraction, calcium handling and sarcomeric organization.^6,24–26^ Similarly, G3BP2 promotes hypertrophy, as its overexpression enhances hypertrophic responses while knockdown attenuates isoproterenol-induced hypertrophy in vivo.^27^ Pathogenic RNA-binding protein condensates have also been described in dilated cardiomyopathy, where the mutant RBM20^R636S^ forms granule arrays along myofibril z-discs in patient myocardium and induced pluripotent stem cell-derived cardiomyocyte (iPSC-CMs). This established the z-disc as a permissive site for biomolecular condensate assembly in the diseased heart.^28^ In acute cardiac disease, SG assembly has been observed in atherosclerosis and myocardial infarction^29–31^, and SG-mediated cardioprotection has been demonstrated in models of atrial fibrillation and sepsis-induced contractile dysfunction.^32,33^ Despite this growing body of evidence, key gaps remain. Previous studies have not determined the precise subcellular localization of SGs within cardiomyocytes, nor have they distinguished the functional consequences of small, newly formed granules from those of larger, persistent assemblies. As a result, the subcellular basis of SG-mediated cardiac pathology remains unresolved.

The cardiomyocyte is a highly polarized cell whose contractile and electrical function depends on the precise spatial organization of its translational and mechanical machinery. In adult cardiomyocytes, ribosomes are not uniformly distributed but instead concentrate at z-disc hotspots and enriched at the intercalated disc.^34–36^ The non-random ribosome distribution is actively maintained by the microtubule motor kinesin-1, which traffics mRNAs and ribosomes along microtubules to discrete subcellular domains, enabling local translation and on-site assembly of contractile components.^37^ Because SG nucleation is initiated by stalled ribosomal complexes, preferential SG assembly at these ribosome-rich domains is a predictable consequence of the translational geography of the cardiomyocyte. The z-disc anchors sarcomeric organization through alpha-actinin, serves as the attachment site for the cardiac dyad, the meeting point at which voltage-sensing L-type calcium channels (Cav1.2) on the t-tubule are positioned in close nanoscale apposition to ryanodine receptors on the junctional sarcoplasmic reticulum to mediate calcium-induced calcium release, which is required for myofilament contraction.^38^ Delivery of Cav1.2 to the cardiac dyad depends on bridging integrator 1 (BIN1)-mediated microtubule-dependent trafficking^39^, placing it in direct competition with any process that disrupts microtubule dynamics or cargo transport. The functional stakes are high: Cav1.2-mediated ICa,L sustains the plateau phase of the ventricular action potential (AP), and even modest reductions in functional channel density shorten AP duration and increase arrhythmia susceptibility.^40,41^ Consistent with this sensitivity, t-tubule loss and Cav1.2 nanodomain disorganization are established features of heart failure that precede overt contractile dysfunction.^42–44^ At the intercalated disc, connexin 43 lateralization and remodeling, hallmarks of arrhythmogenesis, arise from localized oxidative stress concentrated.^45–47^ Supporting the vulnerability of these subcellular domains, pathogenic condensates have been observed in dilated cardiomyopathy, where mutant RBM20^R636S^ forms granule arrays along myofibrillar Z-discs in patient myocardium and iPSC-derived cardiomyocytes.^28^ Together, these observations establish the z-disc and intercalated disc as sites of pre-existing structural vulnerability where SG accumulation, particularly in its coarsened and persistent form, may disproportionately disrupt calcium handling, AP duration, and intercellular coupling.

Despite accumulating evidence implicating SGs in cardiac pathology, three fundamental questions have remained unresolved. First, whether SGs in cardiomyocytes preferentially assemble at mechanically critical sites, such as the z-disc and intercalated disc, rather than distributing generically throughout the cytoplasm. Second, whether the transition to coarsened pathological SGs, particularly within z-disc nanodomains, impairs cardiac electrophysiology. Third, whether SG regulation can be therapeutically targeted to mitigate proarrhythmic remodeling. To address these gaps, SG localization and dynamics were examined under both chronic and acute stress across complementary model systems, and electrophysiological consequences were assessed using pharmacologic perturbations of SG assembly, microtubule integrity, and calcium channel function. These experiments establish that SGs preferentially localize to z-lines and intercalated discs, that SG coarsening, rather than initial formation, represents the critical pathological inflection point driving structural disruption and AP duration shortening, and that coarsened SGs disrupt Cav1.2 nanodomain organization.

## Results

### Stress granules (SGs) localize to mechanically critical sites in chronic disease

SG formation is associated with numerous cardiac pathologies, including atherosclerosis, myocardial infarction, coronary heart disease, and atrial fibrillation.^48–52^ Despite their established role in the cardiac stress response, remarkably little is known about where SGs physically reside within the cardiomyocyte.^32^ The subcellular address of a SG determines which mRNAs it sequesters, which proteins it scaffolds, and ultimately which cellular functions it disrupts.^32^ To address this gap, we investigated SG formation and localization in a controlled system of chronic cardiac disease using the hypertrophic cardiomyopathy mouse model (**Figure 1**) and acute oxidative stress response using sodium arsenite (**Figure 2-4**). In this model, hypertrophic cardiomyopathy was induced by overexpression of a constitutively activated fibroblast growth factor receptor 1 (caFGFR1) in cardiomyocytes in 16-20 week-old mice (**Figure 1A**).^53^

**Figure 1.**
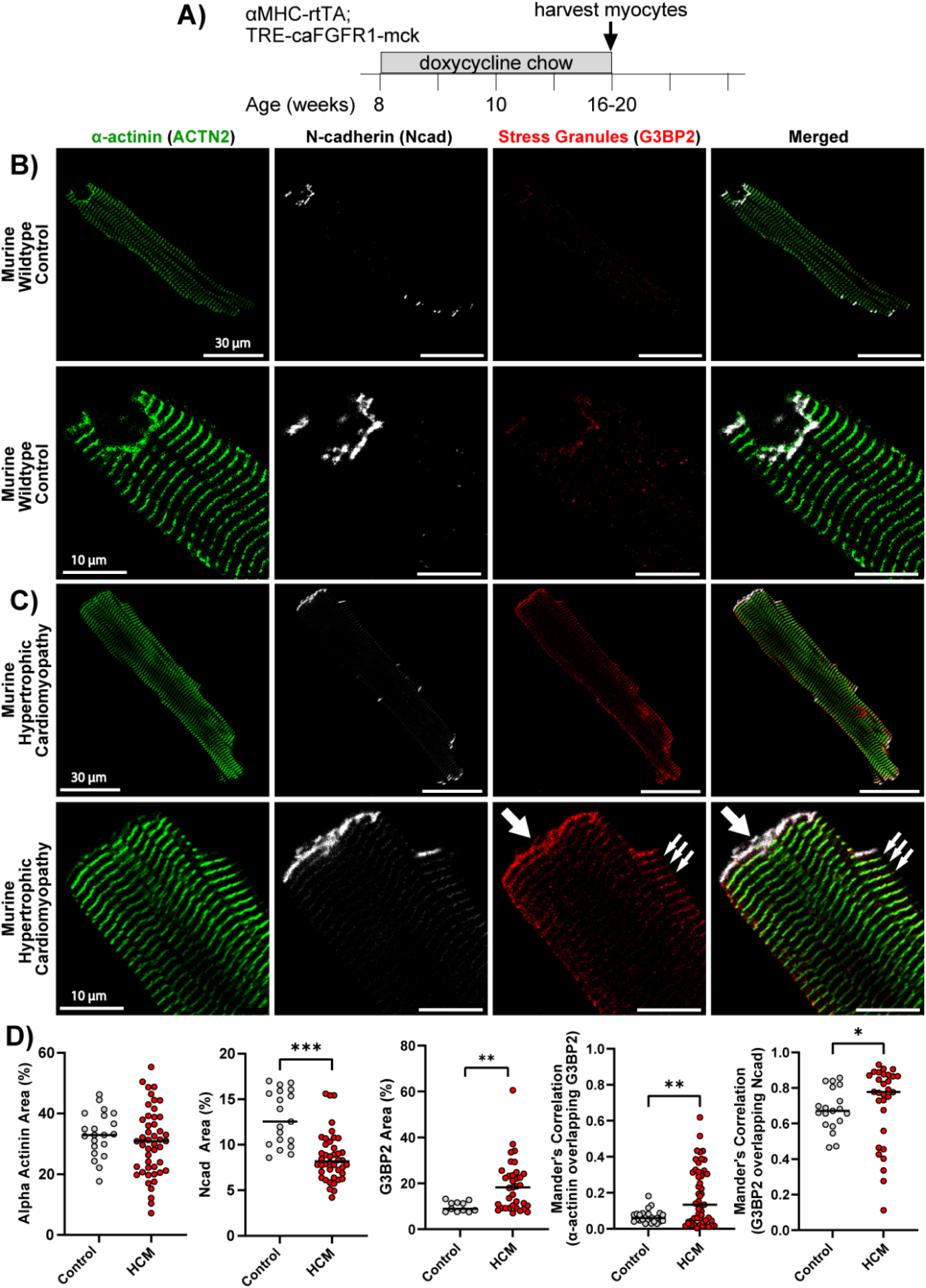
Stress Granules in Chronic Cardiac Disease. **A)** Hypertrophic cardiomyopathy mouse model experimental design. Representative confocal images of isolated cardiomyocytes from **B)** wild-type and **C)** mice with hypertrophic cardiomyopathy (HCM). **B-C)** Isolated cardiomyocytes were stained for z-lines (α-actinin, green) and mechanical junctions known as adherens junctions (N-cadherin, white) to track structural remodeling, as well as G3BP Stress Granule Assembly Factor 2 (G3BP2, red) for stress granule localization. **D)** Structural remodeling was quantified as the percentage area of α-actinin and N-cadherin. Stress granule (SG) formation was quantified by G3BP2 area (%) and Manders’ coefficient for α-actinin and N-cadherin overlap with G3BP2. Murine studies consisted of 3 mice with n = 20 cells for control and n = 45 cells for HCM. Differences in overall distributions were assessed by the Kolmogorov-Smirnov test (NS = P > 0.05; *P ≤ 0.05; **P ≤ 0.01; ***P ≤ 0.001; ****P ≤ 0.0001).

**Figure 2.**
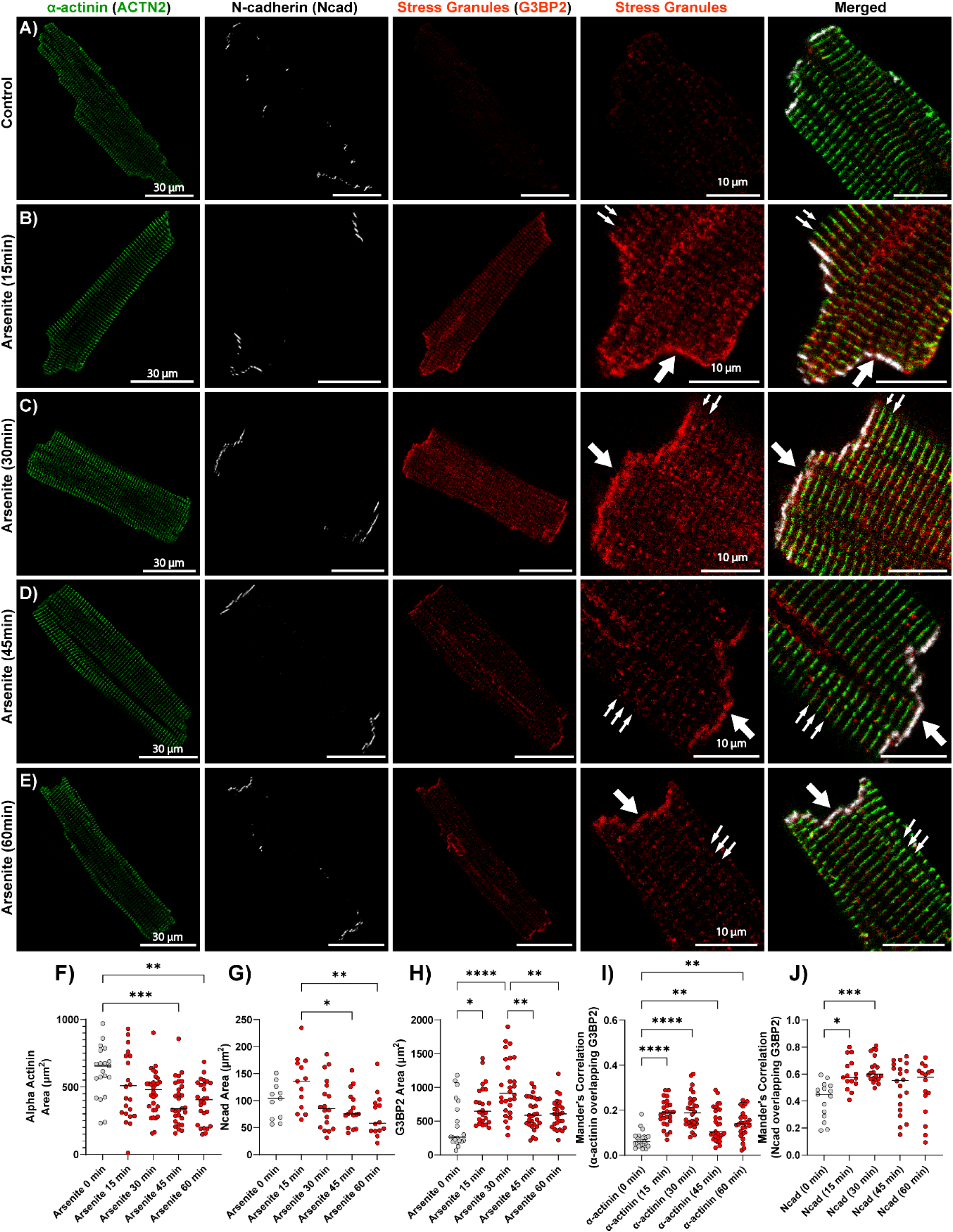
Acute Oxidative Stress in Isolated Murine Cardiomyocytes. Representative confocal images of isolated cardiomyocytes from wild-type mice treated with acute oxidative stress (arsenite 0.5 mM) for 0, 15, 30, 45, and 60 min. **A-E)** Isolated cardiomyocytes were stained for z-lines (α-actinin, green) and mechanical junctions known as adherens junctions (N-cadherin, white) to track structural remodeling, as well as G3BP Stress Granule Assembly Factor 2 (G3BP2, red) for stress granule localization. **F-G)** Structural remodeling was quantified as area (µm^2^) of α-actinin and N-cadherin. Stress granule (SG) formation was quantified by **H)** G3BP2 area (%), **I)** Manders’ coefficient for α-actinin overlap with G3BP2, and **J)** Manders’ coefficient for N-cadherin overlap with G3BP2. Murine studies consisted of 3 biological replicates with n ≥ 20 cells per group. Differences in means were assessed by one-way ANOVA (NS = P > 0.05; *P ≤ 0.05; **P ≤ 0.01; ***P ≤ 0.001; ****P ≤ 0.0001).

SGs were visualized with G3BP2 (red), z-lines with α-actinin (ACTN2, green), and adherens junctions at cell-to-cell ends with n-cadherin (Ncad, white) with confocal microscopy (**Figure 1B-C**). HCM cardiomyocytes showed disrupted mechanical junctions with a reduction in n-cadherin area compared to control cardiomyocytes, while no difference was observed in α-actinin area (**Figure 1D**), indicating that intercalated disc integrity is compromised before sarcomeric structure is affected. SGs nucleate from stalled polysomes and pre-initiation complexes.^54–57^ Because cardiomyocytes concentrate their translational machinery at z-disc hotspots, SG assembly at these landmarks is predicted.^34–36^ HCM cardiomyocytes confirmed this, displaying elevated SG area preferentially localized to z-lines and mechanical junctions (white arrows) rather than dispersed throughout the cytoplasm as previously reported^32^, placing the SG lifecycle in immediate proximity to the sarcomeric and junctional infrastructure on which cardiomyocyte function depends (**Figure 1D**). Next, we set out to further dissect the temporal dynamics of SG formation in an acute oxidative stress model.

### Stress granule coarsening correlates with the disruption of microtubule and α-actinin networks during acute oxidative stress

Sodium arsenite is a well-established agent for studying oxidative stress-induced SG formation. Arsenite-induced oxidative stress was applied to isolated murine wild-type cardiomyocytes (**Figure 2**), whole murine hearts (**Supplementary Figure 1**) and wild-type iPSC-CMs (**Figure 3**). We applied arsenite stress across three complementary model systems using the same dose (0.5 mM), time course (15-60 mins), and marker (G3BP family) previously used to characterize SG nucleation, maturation, and coarsening in non-muscle cells.^54,55,58^ Isolated adult murine cardiomyocytes permit full-thickness confocal imaging at these field-standard timings (**Figure 2**). Whole-heart preparations preserve contractile function during stress but, owing to tissue-level diffusion, require extended exposure of 1 to 2 hrs to achieve comparable SG induction (**Supplementary Figure 1**). Wild-type iPSC-CMs provide a beating, human-protein-composition model at the field-standard timings (**Figure 3**). Convergent findings across all three preparations establish that the SG dynamics described in non-muscle cells operate in the cardiomyocyte and produce cardiomyocyte-specific structural consequences not captured by existing condensate models. Isolated cardiomyocytes and iPSC-CMs were treated with 0.5 mM arsenite for 15, 30, 45, or 60 mins; whole-heart preparations required 1 to 2 hrs. SGs (red), z-lines (green), adherens junctions (white), and microtubules (α-tubulin, white, iPSC-CMs only) were visualized as previously described (**Figures 2A-E, 3A-F**).

**Figure 3.**
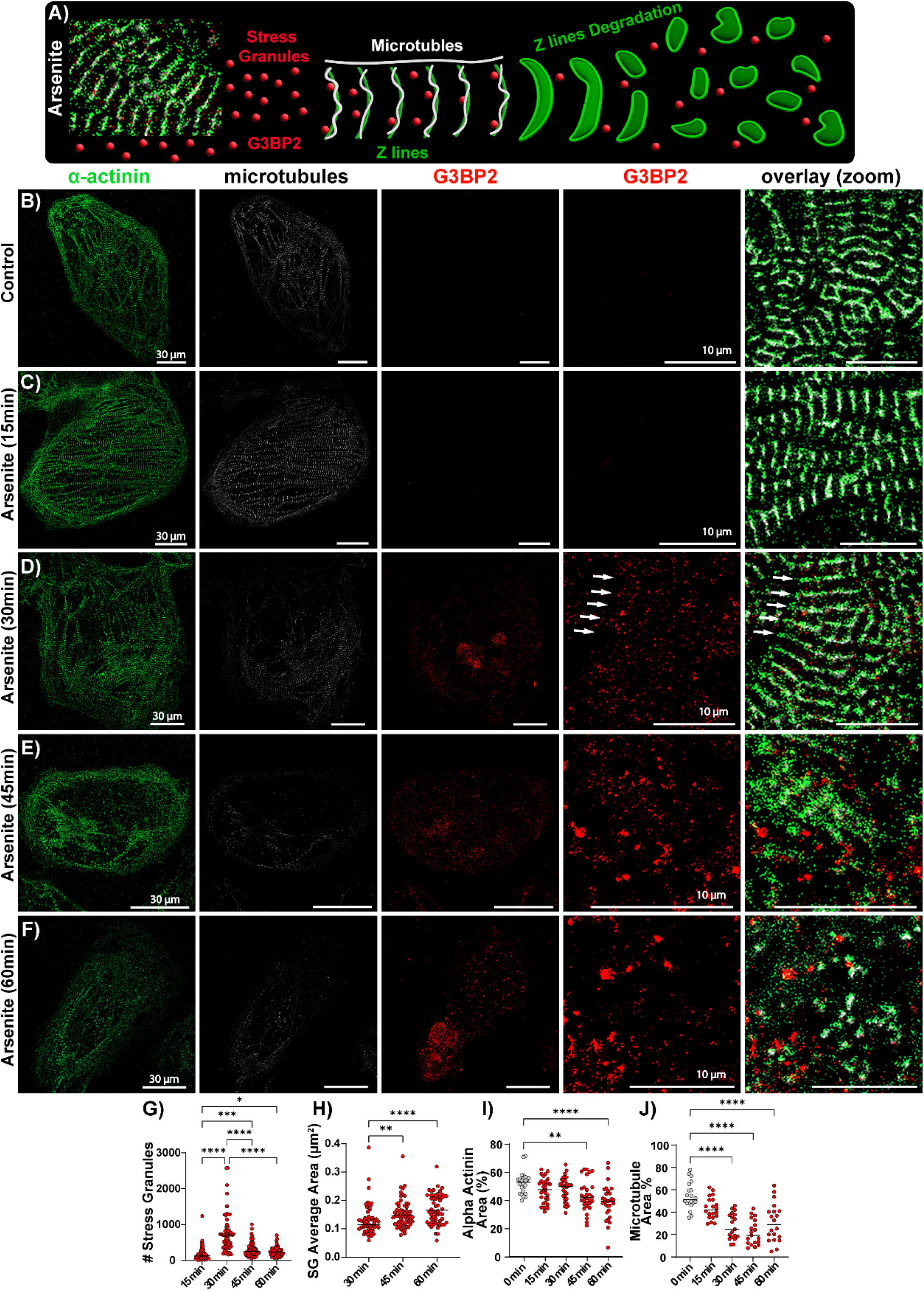
Acute Oxidative Stress in iPSC-CMs. **A)** The cartoon illustrates the close proximity of stress granules (red) to z-lines (green) and microtubules (white), depicting unstressed cardiomyocytes (left), the initial appearance of SGs during acute oxidative stress (30 min arsenite, middle), and progression into structural degradation at longer exposures (45-60 min arsenite, right). **B-F)** Representative confocal images of iPSC-CMs treated with acute oxidative stress (arsenite 0.5 mM) for 0, 15, 30, 45, and 60 min. iPSC-CMs were stained for z-lines (α-actinin, green) and microtubules (α-tubulin, white) to track structural remodeling, as well as G3BP Stress Granule Assembly Factor 2 (G3BP2, red) for stress granule formation. Single plane images are presented as whole cell (30 µm scale bar) and zoomed (5 µm scale bar) perspectives. SG formation was quantified by **G)** number of SGs and **H)** average SG area (n = 30 from 3 replicates). Structural remodeling was quantified by **I)** percentage of α-actinin area and **J)** percentage of microtubule area (n = 30 from 2 replicates). Differences in means were assessed by one-way ANOVA (NS = P > 0.05; *P ≤ 0.05; **P ≤ 0.01; ***P ≤ 0.001; ****P ≤ 0.0001).

**Figure 4.**
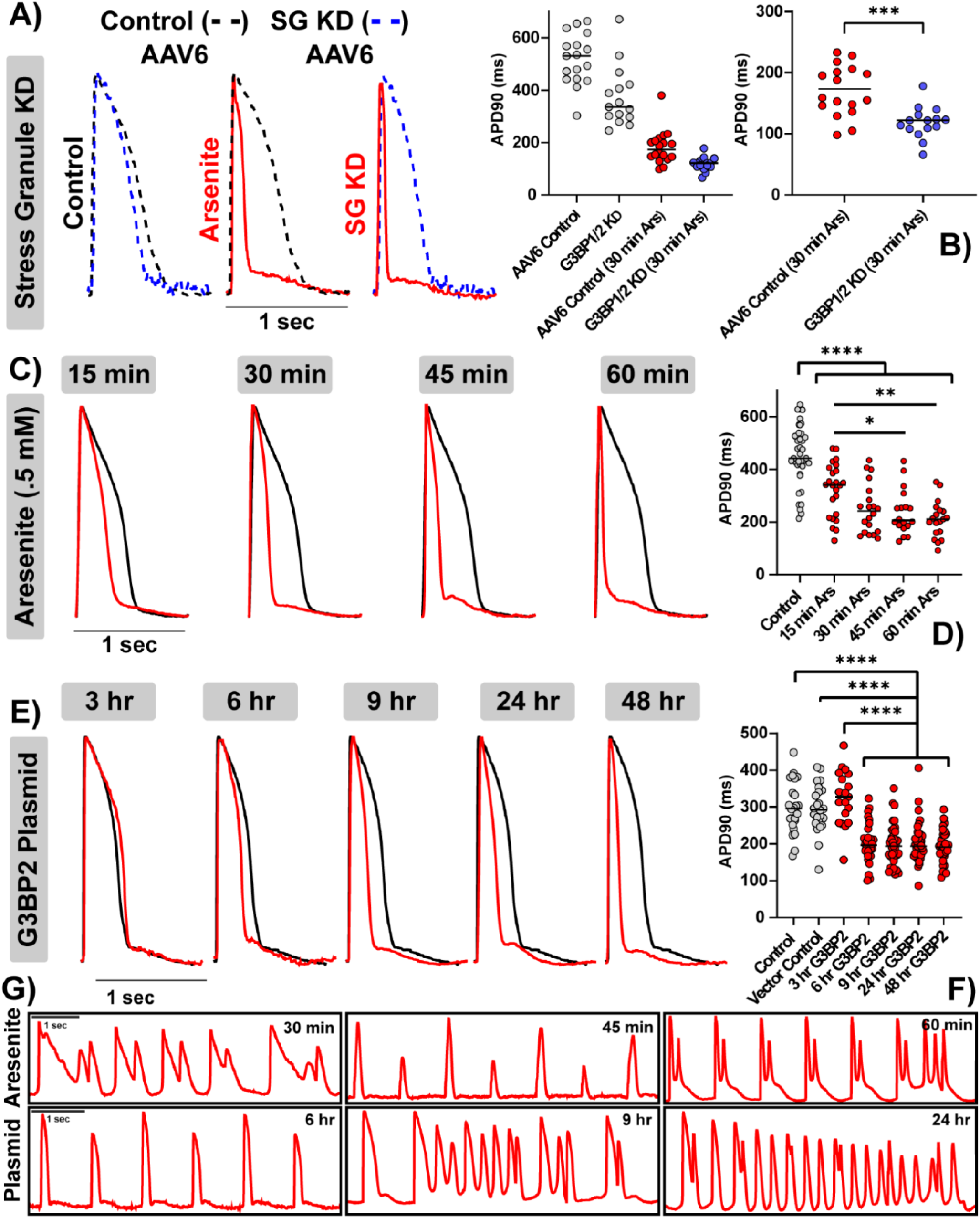
Impact of Stress Granules on Action Potential Morphology and Arrhythmogenesis. Representative action potential (AP) traces from iPSC-CMs subjected to **A)** stress granule (SG) knockdown (KD), **C)** acute oxidative stress (arsenite 0.5 mM), and **E)** SG upregulation (G3BP2 plasmid). **A)** G3BP1 and G3BP2 were knocked down using AAV6 viruses, then cells were challenged with arsenite for 30 min (G3BP1/2 KD (30 min Ars), red right trace). SG KD was compared to the following: empty AAV6 control vector (black dashed trace), SG KD without oxidative stress (G3BP1/2 KD, blue dashed trace), and AAV6 control vector under oxidative stress (AAV6 control (30 min Ars), red middle trace). **C)** Oxidative stress was induced over time at 15, 30, 45, and 60 min with 0.5 mM arsenite (red traces) and compared to untreated iPSC-CMs (black trace). **E)** SG upregulation was induced with a G3BP2 plasmid via magnetofection at 3, 6, 9, 24, and 48 hrs (red traces). SG upregulation was compared to a GFP vector control and an untreated control group (black trace). **G)** Representative AP traces of proarrhythmic phenotypes in iPSC-CMs treated with 0.5 mM arsenite (top row, 30, 45 and 60 min) and SG upregulation (bottom row, 6, 9, and 24 hrs). **B, D, F)** APs were quantified based on the action potential duration at 90% repolarization (APD90). Data was collected from n = 30 recordings across 3 biological replicates. Differences in means were assessed by one-way ANOVA (NS = P > 0.05; *P ≤ 0.05; **P ≤ 0.01; ***P ≤ 0.001; ****P ≤ 0.0001).

In isolated murine cardiomyocytes, SG formation peaked at 30 min of arsenite exposure and declined at 45 and 60 min (**Figure 2H**). Structural consequences emerged only after SG formation peaked: α-actinin and n-cadherin area were preserved at 15 and 30 mins but reduced at 45 and 60 mins (**Figure 2F-G**), placing sarcomeric and mechanical junction disruption downstream of SG assembly rather than concurrent with it. SGs overlapped with both the z-disc and mechanical junctions throughout, with stronger overlap at mechanical junctions (white arrows, **Figure 2I-J**), reproducing the spatial preference observed in the chronic HCM model. Whole-heart preparations recapitulated this sequence, with peak SG formation followed by decline and structural degradation at later timepoints, but required 1 to 2 hrs for comparable SG induction (**Supplementary Figure 1D-E**). iPSC-CMs provided direct quantitative evidence of coarsening: SG number peaked at 30 min and declined at 45 and 60 mins while granule area increased (white arrows, **Figure 3G-H**), indicating that smaller granules were merging into larger ones, a process dependent on microtubule-associated dynein and kinesin motors characterized in non-muscle cells.^13,17,58–60^ As in isolated cardiomyocytes, structural disruption followed rather than accompanied coarsening: α-actinin area was preserved through 30 mins and reduced at 45 and 60 mins (**Figure 3I**), with microtubule networks similarly disrupted at later timepoints (**Figure 3J**).

These data establish that sarcomeric and microtubule disruption follow, rather than accompany, the coarsening transition. Peak SG formation at 30 mins preceded by any structural change, ruling out formation itself as the pathogenic event. Instead, smaller SGs appear to serve as protective translational hubs, with the transition to larger, coarsened granules accompanying pathological consequences.^61,62^ In non-muscle cells, small SGs are resolved through chaperone-mediated pathways, while coarsened granules recruit autophagy-and proteasome-dependent degradation machinery.^14,15,63^ In cardiomyocytes, where structural maintenance depends on continuous protein turnover at the z-disc, sustained degradation activity at coarsened SGs would act directly on the sarcomeric and junctional proteins we observe to be lost.^36,64,65^ Whether coarsened SGs are causally responsible, and whether structural loss carries electrophysiological consequences, is directly tested below.

### Stress granule coarsening, not stress granule formation, drives action potential shortening and arrhythmogenesis

During early stress exposure, SGs halt translation and direct cellular processes to evading cell death. Consistent with a protective role, SGs guard against oxidative stress, calcium overload, and atrial fibrosis in acute atrial fibrillation^32^, and preserve cardiomyocyte contractility during sepsis-induced dysfunction.^33^ In chronic cardiac pathology, G3BP1 upregulation during pressure overload is sufficient to induce SG assembly, and its knockdown attenuates the hypertrophic transcriptome.^25^ G3BP2 similarly promotes hypertrophy through NF-κB activation^27^, underscoring that both assembly factors are dysregulated during pathological remodeling. The z-disc and intercalated disc anchor T-tubule calcium handling nanodomains, including Cav1.2 and ryanodine receptor clusters, which govern AP morphology and excitation-contraction coupling. Disruption of these structures therefore carries direct electrophysiological consequences. Connexin 43 remodeling, a hallmark of heart failure and arrhythmogenesis, arises from oxidative stress at the intercalated disc^45–47^, where enlarged SGs also accumulate (**Figures 2 and 3**). How enlarged, persistent SGs affect cardiac electrophysiology during pathological remodeling remains unknown.

Voltage optical mapping recordings revealed a correlation between SG coarsening and alterations in the cardiac AP. APs were recorded at 1 Hz pacing and quantified as duration at 90% repolarization (APD90). Three paradigms tested whether SGs directly contribute to AP morphology: SG knockdown (AAV6 G3BP1/2 shRNA), acute oxidative stress (0.5 mM arsenite), and SG upregulation (G3BP2 plasmid) (**Figure 4**). To assess the effect of removing SGs during ongoing oxidative stress, G3BP1 and G3BP2 were knocked down for 24 hrs using AAV6 shRNA vectors, then challenged with 0.5 mM arsenite for 30 minutes. An empty GFP AAV6 vector served as a control for viral delivery. Knockdown of SGs in the presence of oxidative stress further shortened APD90 relative to arsenite treatment alone (**Figure 4A-B**), whereas knockdown in the absence of arsenite revealed only modest baseline changes. These data suggest that smaller SGs, which predominate during early oxidative stress, actively prevent AP shortening, consistent with the protective role previously reported in atrial fibrillation.^32^ To characterize how AP morphology evolves as SGs coarsen, APs were recorded at 15, 30, 45, and 60 mins of continuous 0.5 mM arsenite treatment and compared to untreated iPSC-CMs (**Figure 4C-D**). APD90 was progressively shortened over time, with the greatest reductions at 45 and 60 mins. Proarrhythmic activity emerged at 30, 45, and 60 minutes of oxidative stress (**Figure 4G**). This temporal pattern mirrors the structural coarsening data in **Figure 3**, linking granule coarsening directly to APD shortening and arrhythmia susceptibility.

To confirm that APD shortening reflects SG biology rather than nonspecific cellular decline, SGs were induced independently via G3BP2 plasmid magnetofection and APD90 measured at 3, 6, 9, 24, and 48 hours against GFP vector and untreated controls (**Figure 4E-F**). No APD change was detected at 3 hrs, a timepoint at which structural analysis revealed an abundance of small SGs (**Supplemental Figure 2C-D**). Progressive APD90 shortening emerged at 6 hrs and continued through 48 hrs, recapitulating the oxidative stress time course. Proarrhythmic activity emerged at 6, 9, and 24 hrs after transfection, coinciding with the onset and progression of APD90 shortening (**Figure 4G**). The convergence of these findings across mechanistically distinct conditions demonstrates that SG coarsening, rather than oxidative stress, drives APD shortening. Smaller SGs preserve AP morphology, whereas enlarged, persistent SGs are associated with progressive APD90 shortening and increased arrhythmia susceptibility.

### Microtubule inhibition prevents stress granule coarsening and rescues z-disc networks

SGs form as small cytoplasmic foci transported along microtubules by dynein and kinesin motors, enabling successive fusion into larger assemblies.^14,15,17^ Microtubule inhibition disrupts coalescence, yielding more numerous, smaller granules in place of fewer enlarged ones.^13^ Maintaining SGs in this smaller, dispersed state may preserve cytoskeletal integrity by limiting the mechanical burden concentrated at microtubule and z-disc anchor points, where larger condensates impose sustained physical force during transport and nucleation. Persistent coarsening also carries consequences for proteostasis. As SGs grow and become refractory to dissolution, they accumulate ubiquitin chains and recruit VCP/p97 and the 26S proteasome for clearance.^66^ When granules become too large to clear efficiently, sustained proteasome engagement at coarsening sites may deplete local proteolytic capacity, rendering neighboring z-disc and intercalated disc proteins susceptible to accelerated degradation.^67^ We therefore hypothesized that preventing coarsening would preserve α-actinin network integrity.

iPSC-CMs were pretreated with 3 µM nocodazole for 2 hrs prior to acute oxidative stress challenge at 0, 30, 45, and 60 mins (0.5 mM arsenite, **Figure 5**). SG (red) presence, size, and number were quantified relative to z-disc (green) and microtubule (white) networks as previously described (**Figure 5A-D**). Nocodazole sustained an abundance of smaller granules through 45 and 60 minutes, preventing the decline in granule number seen with arsenite alone (white arrows, **Figure 5E**). Without directional motor activity, granules accumulated at the periphery of z-lines rather than trafficking to fusion sites, as shown in the zoomed panel at 45 minutes. SG size remained significantly smaller than arsenite alone at all timepoints, confirming arrested coarsening (**Figure 5F**). Nocodazole preserved α-actinin network organization at 45 and 60 minutes, time points at which arsenite alone produced marked z-line disruption (**Figure 5G**). Microtubule recovery was not observed, consistent with the continued presence of the inhibitor (**Figure 5H**). Microtubule-dependent transport is required for SG coarsening, and arresting coarsening is sufficient to preserve α-actinin integrity during oxidative stress.

**Figure 5.**
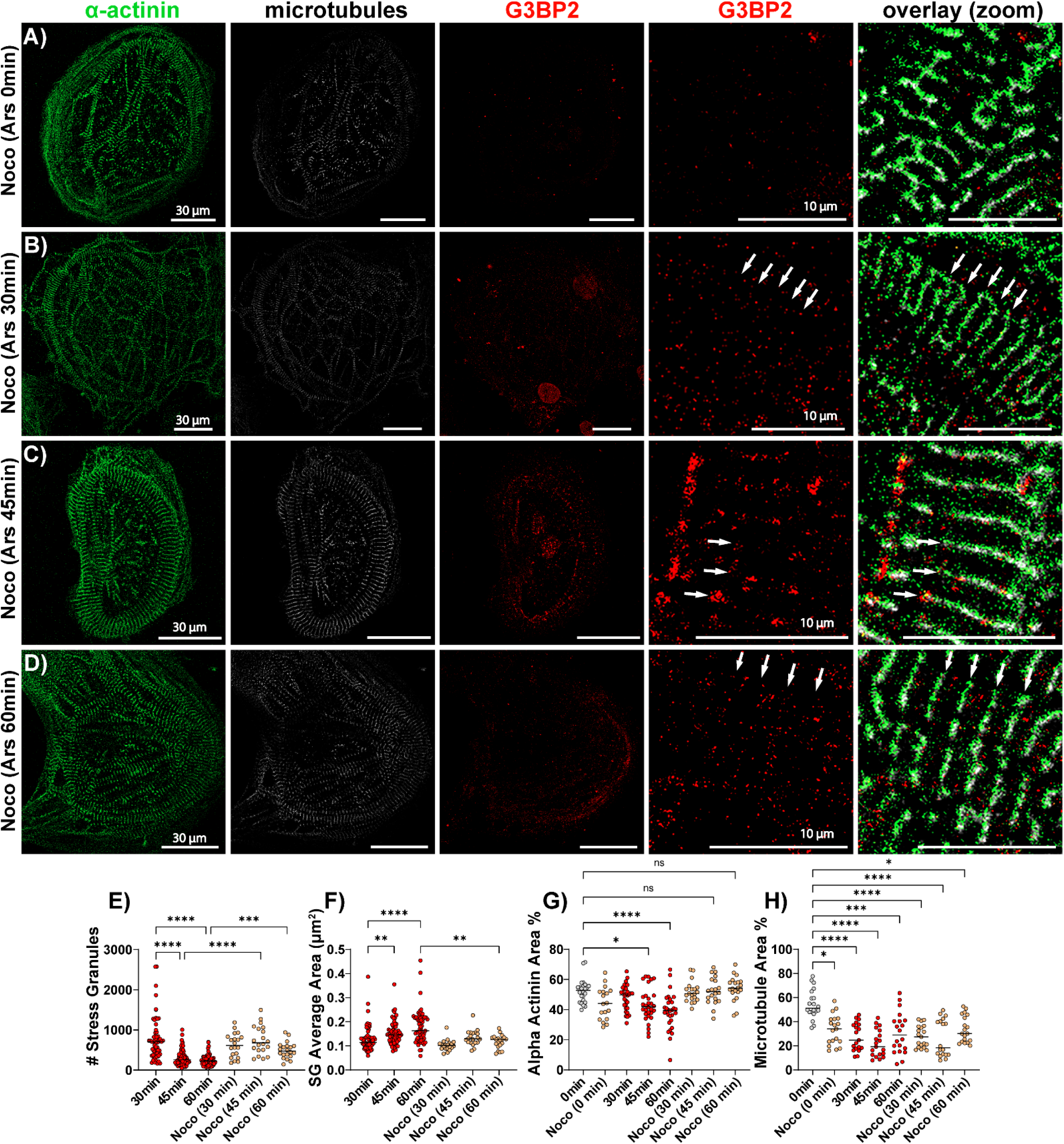
Microtubule Inhibition Prevents Enlargement of Stress Granules and Z-line Degradation. A-D) Representative confocal images of iPSC-CMs pretreated with microtubule inhibitor (nocodazole, Noco, 3 µM), then challenged with acute oxidative stress (arsenite 0.5 mM) for 0, 30, 45, and 60 min. iPSC-CMs were stained for z-lines (α-actinin, green) and microtubules (α-tubulin, white) to track structural remodeling, as well as G3BP Stress Granule Assembly Factor 2 (G3BP2, red) for stress granule (SG) formation. Single plane images are presented as whole cell (30 µm scale bar) and zoomed (5 µm scale bar) perspectives. SG formation was quantified by **E)** number of SGs and **F)** average SG area (n = 20 from 2 replicates). Structural remodeling was quantified by **G)** percentage of α-actinin area and **H)** percentage of microtubule area (n = 20 from 2 replicates). Differences in means were assessed by one-way ANOVA (NS = P > 0.05; *P ≤ 0.05; **P ≤ 0.01; ***P ≤ 0.001; ****P ≤ 0.0001).

*Microtubule inhibition protects calcium handling nanodomains and AP duration*.

The plateau phase of the cardiac AP is sustained by the balance of inward calcium current through L-type calcium channels (Cav1.2) and outward potassium current. Disruption of Cav1.2 nanodomains at T-tubules reduces inward calcium current during the plateau phase, shortening APD90. Given Cav1.2 proximity to disrupted α-actinin networks and prior evidence that nocodazole protects T-tubules^68^, we used confocal microscopy to examine Cav1.2 nanodomain integrity under the same conditions used to map SG coarsening. iPSC-CMs were labeled for G3BP2 (red) and Cav1.2 (yellow) under acute oxidative stress with and without nocodazole pretreatment (**Figure 6**). Oxidative stress reduced Cav1.2 periodicity and area at 45 and 60 minutes (white arrows, **Figure 6A-D, I**), paralleling both APD90 shortening and α-actinin disruption at those timepoints. Strong colocalization between SGs and Cav1.2 (Manders’ coefficient, **Figure 6J**) indicates that coarsening granules accumulate in direct proximity to the calcium handling machinery they appear to destabilize. Nocodazole preserved Cav1.2 periodicity and area (white arrows, **Figure 6E-H, I**), recapitulating α-actinin rescue and suggesting that coarsening consequences converge on Cav1.2 nanodomains. SG colocalization with Cav1.2 was maintained following nocodazole treatment (**Figure 6J**), indicating that proximity to the channel is independent of granule size; it is the coarsening state that determines whether nanodomain integrity is preserved or compromised. SG coarsening destabilizes Cav1.2 nanodomains, providing a structural basis for AP shortening, and arresting coarsening with nocodazole preserves channel organization.

**Figure 6.**
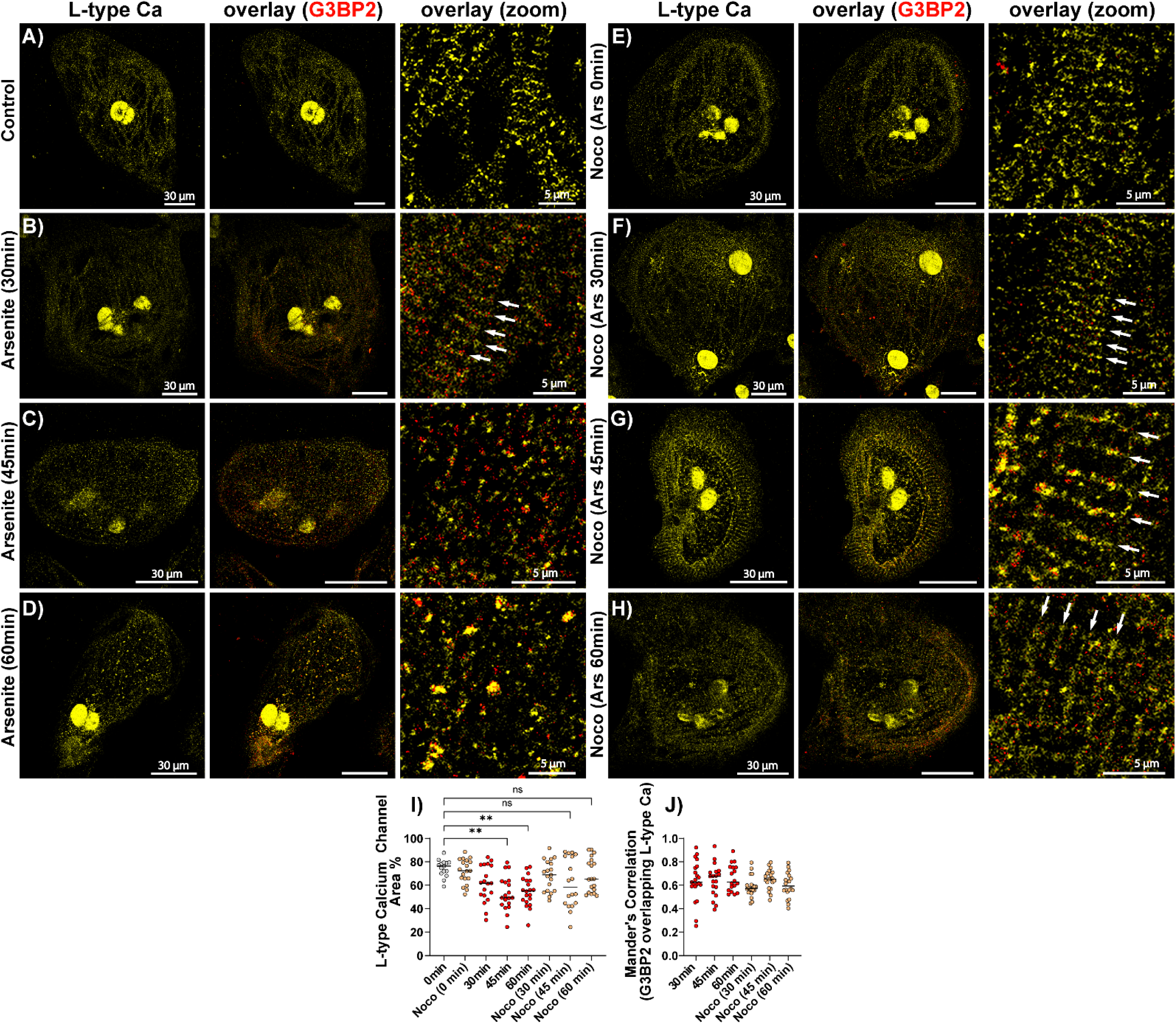
Microtubule Inhibition Prevents L-type Calcium Channel Degradation. Representative confocal images of iPSC-CMs undergoing **A-D)** acute oxidative stress (arsenite 0.5 mM, 0, 30, 45, and 60 min) and **E-H)** pretreatment with microtubule inhibitor (nocodazole, Noco, 3 µM) followed by acute oxidative stress (0.5 mM arsenite) for 0, 30, 45, and 60 min. iPSC-CMs were stained for L-type calcium channels (Cav1.2, yellow) to track structural remodeling within the calcium handling machinery, as well as G3BP Stress Granule Assembly Factor 2 (G3BP2, red) for stress granule association. Single plane images are presented as whole cell (30 µm scale bar) and zoomed (5 µm scale bar) perspectives. L-type calcium channels were quantified by **I)** percentage Cav1.2 area and **J)** Manders’ coefficient for G3BP2 overlap with L-type calcium channels (n = 20 from 2 replicates). Differences in means were assessed by one-way ANOVA (NS = P > 0.05; *P ≤ 0.05; **P ≤ 0.01; ***P ≤ 0.001; ****P ≤ 0.0001).

Prior work demonstrated that nocodazole increases calcium transient amplitude and prevents spontaneous calcium release predicting that microtubule inhibition would protect against oxidative stress-induced AP shortening.^68^ To test this, iPSC-CMs were pretreated with 3 µM nocodazole^13^ for 2 hrs prior to 0.5 mM arsenite challenge, and APs were recorded at 15, 30, 45, and 60 mins alongside arsenite alone (red traces) and untreated (dashed black trace), DMSO (black trace), and nocodazole-only controls (tan traces, **Figure 7A**). AP duration was quantified at 80% repolarization (APD80). APD80 repolarization was used due to altered morphology during phase 4 of the AP. Arsenite (red dots) produced progressive APD80 shortening from 30 to 60 minutes relative to all controls (**Figure 7B**), consistent with the time-dependent coarsening and Cav1.2 disruption described above. Nocodazole (tan dots) abolished this shortening, with APD80 values at 30 and 45 minutes remaining not significantly changed from untreated controls (**Figure 7B**). These findings connect Cav1.2 nanodomain preservation in **Figure 6** to a functional electrophysiological outcome, strengthening the conclusion that microtubule-driven coarsening underlies APD shortening and arrhythmia susceptibility.

**Figure 7.**
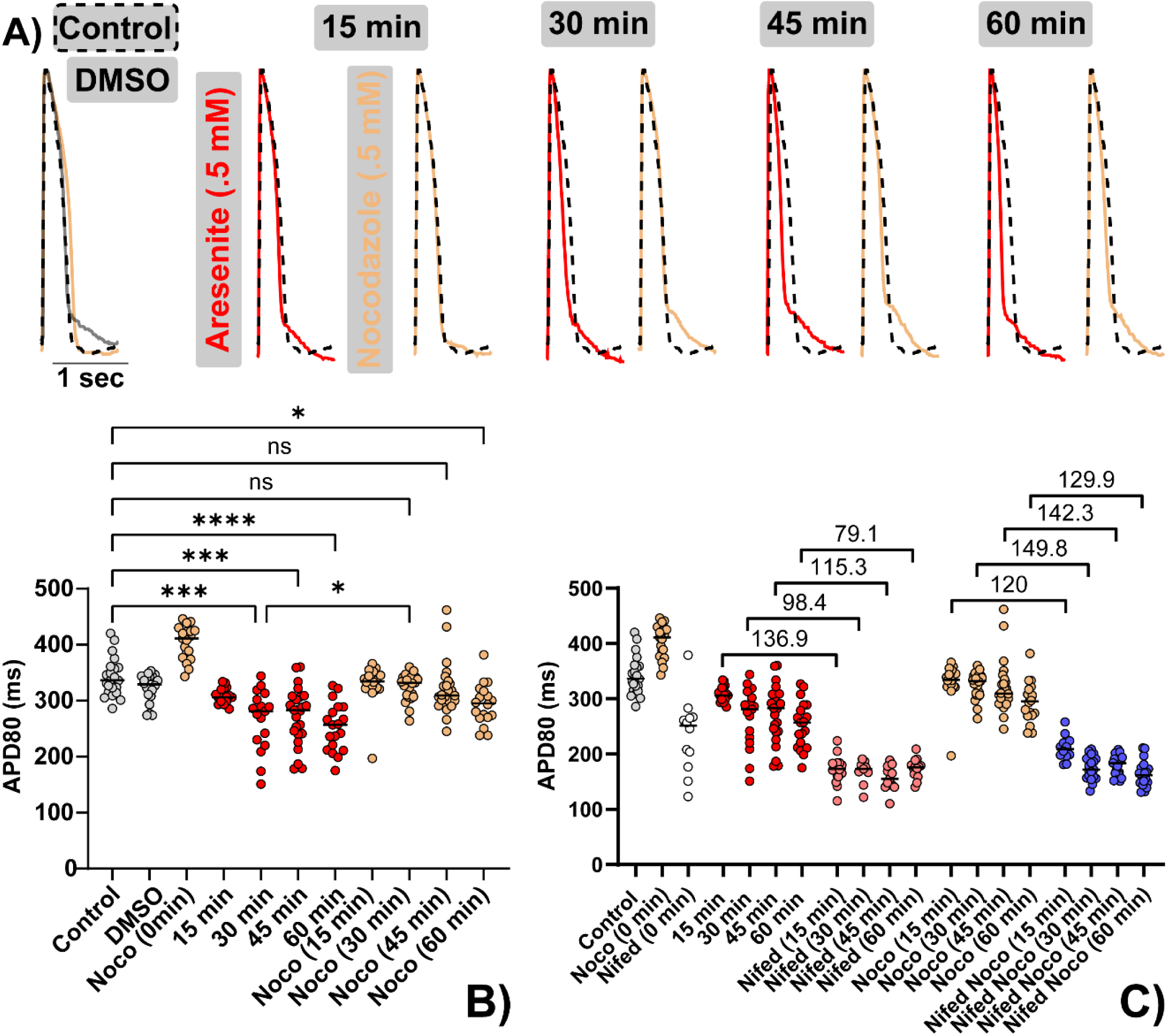
Microtubule Inhibition Prevents Action Potential Shortening. **A)** Representative action potential (AP) traces of iPSC-CMs pretreated with **B)** microtubule inhibitor (nocodazole, Noco, 3 µM), then challenged with acute oxidative stress (arsenite 0.5 mM) for 0, 15, 30, 45, and 60 min (tan traces). Nocodazole pretreatment was compared with acute oxidative stress alone (red traces), an untreated control (dashed black trace), a DMSO vehicle control (black trace), and nocodazole without oxidative stress (Noco 0 min, tan trace). APs were quantified based on the action potential duration at 80% repolarization (APD80). **C)** A calcium channel blocker (nifedipine, 25 nM) was applied acutely to assess the contribution of L-type calcium channel current to APD under acute oxidative stress (pink) compared to nocodazole pretreatment with acute oxidative stress (blue). The difference between medians was calculated to quantify APD reduction following acute nifedipine treatment. Data was collected from n = 30 recordings across 3 biological replicates. Differences in means were assessed by one-way ANOVA (NS = P > 0.05; *P ≤ 0.05; **P ≤ 0.01; ***P ≤ 0.001; ****P ≤ 0.0001).

To determine whether nocodazole’s APD80 protection depends on preserved Cav1.2 current, nifedipine (25 nM) was applied acutely before each timepoint during optical mapping with and without nocodazole pretreatment (**Figure 7C**). If nocodazole acts by sustaining functional Cav1.2 channels, then pharmacological block of those channels should selectively diminish the protective effect and converge APD80 values across treatment groups regardless of timepoint. This prediction was confirmed. Nifedipine reduced APD80 to a comparable floor across all timepoints and conditions, indicating that the residual calcium current remaining after oxidative stress and the larger calcium current preserved by nocodazole are both carried through Cav1.2. The greater absolute reduction in the nocodazole-pretreated group (blue dots) relative to oxidative stress alone (pink dots) reflects the higher functional channel reserve maintained by microtubule inhibition: because nocodazole protected more Cav1.2 channels from oxidative stress-induced disruption, nifedipine had proportionally more current to block. Together with **Figure 6**, these findings establish that microtubule inhibition prevents AP shortening by sustaining Cav1.2 current through structural protection of channel organization.

## Discussion

### Stress granule localization to mechanically critical sites redefines the cardiac stress granule paradigm

The present study demonstrates that SGs in cardiomyocytes are not generic cytoplasmic condensates but site-specific structures that preferentially assemble at mechanically critical nanodomains, including the z-disc and the intercalated disc. Prior cardiac studies have described SGs as large particles scattered throughout the cytoplasm following tachypacing^32^, distributed without reference to subcellular landmarks in sepsis-challenged cardiomyocytes^33^, or accumulating in a perinuclear cytoplasmic pattern upon G3BP1 overexpression.^25^ The present findings contrast with these accounts, identifying the z-disc and intercalated disc as preferred assembly sites. A mechanistic basis for this localization is provided by the organization of the cardiac translational machinery: ribosomes in adult cardiomyocytes are concentrated at the z-disc and enriched at the intercalated disc^34–36^, with microtubule-dependent kinesin transport required to maintain their localization at these sites.^37^ Because SG nucleation is initiated by eIF2α phosphorylation and the subsequent recruitment of stalled 40S ribosomal subunits and preinitiation complexes into condensate seeds^56,69,70^, their preferential assembly at z-disc and intercalated disc hotspots is a predictable consequence of the cardiomyocyte’s ribosome geography.

Precedent for RNP condensate localization to cardiac z-discs exists in the pathogenic granules formed by mutant RBM20 in dilated cardiomyopathy, where sarcoplasmic RBM20R636S decorates myofibril z-discs in a rosary-like configuration^28^, reinforcing the z-disc as a permissive site for biomolecular condensate assembly in the diseased heart. In the HCM model, SG accumulation at mechanical junctions coincided with n-cadherin loss at the intercalated disc, consistent with reported reductions in cadherin-catenin components across cardiomyopathy explants^71^, suggesting that chronic stress drives SG residence at sites of pre-existing structural vulnerability. A higher degree of colocalization between G3BP2-marked SGs and the cardiac structural landmarks α-actinin and n-cadherin was observed across both chronic (HCM mouse model) and acute stress (arsenite induced oxidative stress) contexts, encompassing isolated murine cardiomyocytes, whole-heart preparations, and iPSC-CMs. Such spatial specificity places the SG lifecycle in direct proximity to the sarcomeric and mechanical junction infrastructure on which cardiomyocyte function depends and indicates that subcellular address determines which mRNAs are sequestered and which structural nanodomains are exposed to granule-associated degradation machinery.

### The coarsening transition, not stress granule formation, is the pathological inflection point

A second central finding is that the coarsening transition, not initial SG formation, is the event that drives structural and electrophysiological dysfunction. Prior cardiac studies have established a broadly protective role for SG formation: inhibition of SG assembly worsened LPS-induced cardiomyocyte dysfunction^33^, and G3BP1-mediated SG induction attenuated oxidative stress, calcium overload, and atrial fibrosis in a tachypacing model of atrial fibrillation.^32^ However, neither study distinguished between the functional consequences of small, newly assembled granules and coarsened, persistent ones, a distinction the present data resolve. Peak SG abundance at 30 minutes, when granules are most numerous and smallest, coincided with preserved structural integrity; knockdown at this timepoint further shortened action potential duration (APD), confirming active cardioprotection by small SGs. At 45 and 60 minutes, granule number declined while total granule area increased, marking the coarsening transition, after which α-actinin, n-cadherin, and microtubule networks were disrupted and APD shortening and proarrhythmic activity emerged. SG upregulation at 3 hours, when small granules predominate, had no effect on AP morphology, while progressive APD shortening emerged at 6 hours and beyond, coinciding with granule coarsening. Together, these data establish that granule size and coarsening state, rather than granule presence, are the critical determinants of pathological outcome.

The mechanistic basis for this transition lies in the differential clearance of small versus coarsened SGs. Under acute stress, SGs disassemble within 60 to 120 minutes upon stress removal through chaperone-mediated pathways involving HSP40, HSP70, and HSP104.^21^ When sustained or chronic stress exhausts these chaperone resources, or when competing protein aggregates divert chaperone availability, SG disassembly fails, granules persist, and the liquid-to-solid transition that underlies pathological granule behavior ensues.^22^ Larger, persistent granules that cannot be resolved by chaperones require escalating clearance: autophagy-dependent degradation and active recruitment of the 26S proteasome and ubiquitin-selective segregase p97 to the granule surface.^9,21^ ZFAND1 mediates this escalation by recruiting both the 26S proteasome and p97 directly to arsenite-induced SGs; in its absence, granules accumulate defective ribosomal products and persist, converting to aberrant, disease-linked structures.^23^ In cardiomyocytes, this escalating clearance machinery is recruited to coarsened granules at the z-disc and intercalated disc. The 26S proteasome and p97 may be localized in close proximity to α-actinin, N-cadherin, and microtubule networks. Their selective loss at timepoints coinciding with granule coarsening suggests a mechanistic link between impaired protein quality control, granule persistence, and the structural disruption that precedes electrophysiological dysfunction.

### Cav1.2 nanodomain disruption bridges structural and electrophysiological findings

Cav1.2 channels are concentrated at cardiac T-tubules, invaginations of the sarcolemma that traverse the cell along the z-line and position L-type calcium channels in close apposition to ryanodine receptors on the junctional sarcoplasmic reticulum.^38^ The BIN1 scaffolding protein organizes this nanodomain by directing microtubule-dependent trafficking of Cav1.2 to the T-tubule membrane, such that perturbation of microtubule integrity disorganizes channel delivery.^39^ Through this arrangement, the rapidly activating ICa,L provides the sustained inward current that underlies the plateau phase of the ventricular AP, and even modest reductions in functional channel density shorten APD and increase arrhythmia susceptibility.^40,41^ T-tubule loss and Cav1.2 nanodomain disorganization are recognized features of heart failure across diverse etiologies and, critically, emerge before the onset of overt contractile dysfunction.^42–44^ The present findings identify SG coarsening as an upstream event in this remodeling program. Cav1.2 periodicity and area were reduced at 45 and 60 minutes of acute oxidative stress, temporally coinciding with the onset of APD shortening, and SGs colocalized with Cav1.2 nanodomains throughout the time course. Microtubule inhibition with nocodazole prevented SG coarsening, preserved both z-disc integrity and Cav1.2 nanodomain organization, and abolished APD shortening. SG colocalization with Cav1.2 was maintained after nocodazole treatment, indicating that nanodomain proximity is size-independent and that pathological consequences require granule coarsening rather than granule presence alone. Cav1.2-dependence of this protection was confirmed pharmacologically: nocodazole-pretreated cells exhibited a greater reduction in APD80 upon nifedipine block, consistent with a higher reserve of functional channels preserved by microtubule inhibition. Together, these data establish that microtubule-driven SG coarsening at the z-disc-Cav1.2 nanodomain is the proximate cause of APD shortening and arrhythmia susceptibility in the setting of cardiac oxidative stress.

### Microtubule-dependent stress granule transport as a targetable pathway

The microtubule cytoskeleton is required for SG coarsening. Dynamic microtubule turnover generates long-range granule movements that drive fusion of small granules into large perinuclear assemblies, and disruption of this transport arrests coarsening by leaving granule nucleation intact while eliminating the secondary coalescence step.^13–15^ Consistent with this established mechanism, nocodazole and colchicine prevent SG fusion and maintain granule populations below the pathological size threshold across multiple cell systems.^16,17^ The present findings establish that this pathway operates in cardiomyocytes with direct electrophysiological consequences: nocodazole pretreatment prevented SG coarsening, preserved alpha-actinin network integrity and Cav1.2 nanodomain organization, and abolished APD shortening, defining a causal chain from granule size through structural remodeling to electrophysiological dysfunction. These results are consonant with prior work demonstrating that microtubule depolymerization protects excitation-contraction coupling machinery in cultured cardiomyocytes: nocodazole and colchicine prevented junctophilin-2 redistribution from T-tubule membranes, attenuated T-tubule impairment, increased calcium transient amplitude, and reduced calcium spark frequency.^68,72^ Together, these observations position microtubule-dependent SG transport as a druggable node upstream of both structural and arrhythmogenic remodeling in the setting of oxidative stress.

Although nocodazole lacks the selectivity required for therapeutic application, the functional rescue it produces establishes proof of concept for targeting the coarsening step. The molecular motors responsible for SG transport make distinct contributions: dynein, which drives cargo toward the microtubule minus-end and the perinuclear interior, is required for the initial phases of SG assembly and for the subsequent coalescence that produces large granules, while kinesin participates in SG disassembly and limits further granule growth, such that kinesin depletion leads to persistent SG accumulation.^17–20^ Selective pharmacological manipulation of these motors therefore represents a mechanistically motivated strategy for controlling granule size without eliminating granule formation: future experiments in iPSC-CMs using the kinesin-1 activator kinesore and the dynein inhibitor sodium orthovanadate will define which motor activities drive coarsening in the cardiomyocyte specifically.^73,74^ A broader implication concerns chronic cardiac disease. G3BP1 and G3BP2, the core SG assembly factors studied here, are both upregulated in cardiac hypertrophy^25,27^, raising the possibility that chronic elevation of these nucleating factors primes cardiomyocytes toward a low-threshold coarsening state. Whether this chronic sensitization contributes to the arrhythmia burden observed in HCM and heart failure patients is a question the present work directly motivates.

### Limitations

Several limitations were present. The electrophysiological experiments were performed entirely in iPSC-CMs (WTc-11 line), a model that recapitulates key features of human cardiomyocyte biology but lacks the fully developed T-tubule networks of adult ventricular myocytes. Because Cav1.2 nanodomains are T-tubule-resident structures, conclusions about nanodomain organization should be interpreted in the context of the immature T-tubule architecture characteristic of this system. Prior to the experiments reported here, the presence of sarcomeric z-disc, microtubule networks, and Cav1.2, as well as sodium channels (Nav1.5), ryanodine receptors, and potassium channels (Kir2.1 and Kv7.1) colocalized with alpha-actinin, was confirmed in control iPSC-CMs. Validation in adult primary cardiomyocytes and in vivo arrhythmia models will be required to confirm translation to the intact heart. The chronic model was limited to a single HCM etiology (caFGFR1 overexpression), and whether preferential SG localization to z-lines and intercalated discs generalizes to pressure overload, dilated cardiomyopathy, or post-infarction remodeling remains to be tested. Additionally, while isolated murine cardiomyocytes and iPSC-CMs were challenged with identical arsenite doses and exposure times, whole-heart preparations required longer exposure to achieve comparable SG induction, reflecting tissue-level diffusion barriers. Prior work characterizing cardiac stress responses in this context was conducted predominantly in left ventricular myocardium, with cardiomyocytes isolated from the apical region and functional assessments reflecting primarily left-sided physiology, without dedicated right ventricular histology or functional analysis. In the present study, cardiomyocytes were isolated from whole hearts encompassing both left and right ventricular myocardium, and whether SG coarsening dynamics, CaV1.2 nanodomain disruption, and electrophysiological consequences differ between left and right ventricular cardiomyocytes was not assessed and represents an important direction for future investigation.

The use of sodium arsenite carries the inherent limitation that arsenite exerts pleiotropic cellular effects, including enzyme inhibition, protein misfolding, and direct thiol oxidation, beyond oxidative stress-induced SG formation.^75–77^ Three converging lines of evidence mitigate this concern. First, G3BP2 overexpression induced SG coarsening independently of arsenite and produced progressive APD90 shortening from 6 to 48 hours, demonstrating that SG coarsening, not arsenite exposure, drives the electrophysiological phenotype. Second, G3BP1/G3BP2 knockdown in the continued presence of arsenite further shortened APD90 beyond arsenite alone, confirming that SGs provide protection during early stress. Third, nocodazole pretreatment arrested SG coarsening while oxidative stress continued, and fully rescued both structural integrity and APD duration, establishing that the observed pathology tracks with coarsening state rather than with arsenite exposure alone.

Nocodazole depolymerizes all cytoplasmic microtubules and disrupts vesicle trafficking, protein delivery, and structural maintenance in addition to SG transport; the contribution of these non-SG effects to the observed rescue cannot be fully excluded. Future experiments will address this by targeting the specific motor proteins responsible for SG coalescence. Selective pharmacological manipulation of these activities using the kinesin-1 activator kinesore and the dynein inhibitor sodium orthovanadate in iPSC-CMs will determine which motor functions are responsible for coarsening in the cardiomyocyte and will enable attribution of any functional rescue to SG-specific transport rather than global microtubule depolymerization.^73,74^

## Materials and Methods

### Sex as a Biological Variable

Our study examined male and female mice, and similar findings were reported for both sexes.

### Cell Lines

Wild-type male human induced pluripotent stem cells (hiPSCs; WTC11, Coriell GM25256) were obtained from The Genome Engineering and Stem Cell Center (GESC@MGI) at Washington University in St. Louis School of Medicine. hiPSCs were cultured in mTeSR Plus medium (STEMCELL Technologies) on 6-well plates coated with Matrigel (1:100, Corning) and grown at 37°C and 5% CO_2_. hiPSCs were grown to 80-90% confluency before passaging or differentiation into cardiomyocytes (iPSC-CMs). hiPSCs were passaged as clusters using Versene (Thermo Fisher Scientific) into 6-well plates for hiPSC maintenance or 12-well plates for cardiomyocyte differentiation. Once hiPSCs reached 90% confluency, they were differentiated into cardiomyocytes via a well-established protocol using small molecule manipulation of Wnt signaling.^78,79^ On differentiation day 0, medium was changed to RPMI 1640 (Life Technologies) with B27 without insulin (B27- , 2%, Life Technologies), ascorbic acid (AA, 150 µg/mL, Fisher Scientific), and CHIR99021 (6 µM, Biogem). On day 2, exactly 48 hours after initial CHIR mediated WNT suppression, medium was changed to RPMI 1640 with B27- (2%), AA (150 µg/mL), and IWP2 (2.5 µM, Biogem). On day 4, medium was changed to RPMI 1640 with B27- (2%) and AA (150 µg/mL). Lastly, on day 6, medium was changed to RPMI 1640 with B27 with insulin (2%, Life Technologies) (RPMI-C). Following differentiation, hiPSC derived cardiomyocyte (iPSC-CM) cultures were fed RPMI-C every 2 days. Beating sheets of iPSC-CMs were replated at 1×10^6^ cells/well on day 14 to prevent sheet detachment and death. Off-target cells (non-cardiomyocytes) were removed via lactate purification on days 20 and 22. Lactate purification medium was composed of RPMI 1640 without glucose (Life Technologies) with lactate (4 mM, Sigma-Aldrich), non-essential amino acids (1%, Fisher Scientific), and Glutamax (1%, Fisher Scientific). iPSC-CMs were replated for downstream structural and functional analysis on days 27-30. Cells were dissociated into single cells and replated at 40,000 cells per 35 mm glass-bottom dish for confocal microscopy and 60,000-80,000 cells per plastic-bottom dish for optical mapping. Presence of iPSC-CMs was validated by spontaneous beating and immunostaining for cardiac structural markers (α-actinin).

### Hypertrophic Cardiomyopathy Mouse Model

Hypertrophic cardiomyopathy (HCM) was induced by chronic cardiomyocyte-specific expression of constitutively active fibroblast growth factor receptor 1 (caFGFR1) in mice.^53^ Long-term caFGFR1 overexpression produces an HCM phenotype characterized by concentric hypertrophy, preserved systolic function, and patchy fibrosis with myocyte disarray.^53^ TRE-caFGFR1 mice were generated by cloning FGFR31C(R248C)-c-myc cDNA into the pTRE2 vector; the linearized vector was then injected into fertilized FVB oocytes to establish founder lines. Founders were crossed with αMHC-rtTA mice, and 8 week old double-transgenic offspring were given doxycycline chow for 4 weeks, to induce transgene expression. Hearts were harvested at 16-20 weeks for myocyte isolation. Mice were maintained on a mixed C57BL/6J-129X1/SvJ genetic background, and both sexes were used. All procedures complied with the Guide for the Care and Use of Laboratory Animals (NIH publication No. 85-23, revised 1996), and all protocols were approved by the Animal Studies Committee at Washington University School of Medicine.

### Single Mouse Ventricular Cardiomyocyte Isolation

Adult mice (16–20 weeks) were deeply anesthetized with 2.5% Avertin (0.01 mL/g, IP) and heparinized (5 IU/g, IP) before heart removal. The excised heart was rapidly cannulated via the aorta and mounted on a Langendorff perfusion apparatus, then perfused for 2–3 min with a HEPES-buffered solution (in mM: 140 NaCl, 5.4 KCl, 1.2 MgCl2, 1 CaCl2, 10 HEPES, 10 glucose, 20 taurine, 5 adenosine; 10 IU/mL Na-heparin; pH 7.4) at 4 mL/min. Next, a low Ca²⁺-high K⁺ solution supplemented with EGTA (in mM: 115 NaCl, 14 KCl, 1.2 MgCl2, 0.025 CaCl2, 10 HEPES, 10 glucose, 20 taurine, 5 adenosine, 0.3 EGTA; pH 7.2) was perfused for 6 min, followed by 15–20 min of perfusion with an enzyme solution (in mM: 115 NaCl, 14 KCl, 1.2 MgCl2, 0.025 CaCl2, 10 HEPES, 10 glucose, 20 taurine, 5 adenosine, 1.37 mg/mL collagenase II (295 U/mg, Worthington Biochemical Corp., United States) and 1.3 µg/mL protease XIV, pH 7.4). All solutions were maintained at 36°C and continuously oxygenated. After digestion, ventricles were excised along the atrioventricular border, minced in enzyme solution, gently agitated to disperse cells, and filtered through a nylon mesh. Digestion was stopped by adding a low Ca²⁺-high K⁺ solution supplemented with bovine serum albumin (5 mg/mL), and the cell suspension was centrifuged for 3 min at 23 × g (Eppendorf Centrifuge 5920R, Germany). Ventricular cardiomyocytes were then resuspended in fresh low Ca²⁺-high K⁺ solution and plated on Matrigel-coated (1:100, Corning, United States) glass coverslips, fixed for 5 min at room temperature with 2% paraformaldehyde (PFA) in PBS, washed in PBS (3 × 10 min at room temperature), and stored in PBS at 4°C until immunolabeling.

### Cellular Treatments

*<u>Oxidative Stress:</u>* Sodium arsenite is a commonly used chemical in the condensate field to induce oxidative stress-induced SGs.^80–82^ Acute oxidative stress was induced in iPSC-CMs with 0.5 mM sodium arsenite dissolved in RPMI-C medium. iPSC-CMs were incubated at 37°C and 5% CO₂ for 15, 30, 45, and 60 min. Then, iPSC-CMs were optically mapped or fixed for immunofluorescence confocal imaging.

*<u>Drug Interventions (nocodazole and nifedipine):</u>* A microtubule inhibitor (nocodazole) was used to prevent SG transport along microtubules. iPSC-CMs were pretreated with nocodazole (3 µM)^13^ for 2 hrs before acute oxidative stress challenge (0.5 mM arsenite) at 15, 30, 45, and 60 min. Then, samples were prepared for optical mapping and immunofluorescence staining for confocal microscopy. A calcium channel blocker (nifedipine) was used to determine the contribution of calcium current to the cardiac AP among three groups: control, oxidative stress, and microtubule inhibitor pretreatment prior to oxidative stress. Nifedipine (25 nM)^83^ was added acutely (1–3 min) before recording each timepoint with optical mapping.

*<u>Stress Granule Knockdown:</u>* Studies have shown that knockdown of both G3BP1 and G3BP2 is required for inhibition of SG formation.^84^ SG knockdown (KD) was achieved with adeno-associated virus type 6 (AAV6) containing shRNA targeting G3BP1 (pAAV[shRNA]-mCherry-U6>hG3BP1[shRNA#1], VB900150-4232jcm) and G3BP2 (pAAV[shRNA]-EGFP-U6>hG3BP2[shRNA#3], VB900147-9031shm) from Vectorbuilder. Scrambled shRNA control AAV6 viruses (GFP) served as controls for AAV6 viral transduction. iPSC-CMs were transduced with AAV6 G3BP1 and G3BP2 shRNA in RPMI-C medium for 24 hrs (multiplicity of infection: 30,000) prior to arsenite challenge (30 min). AAV6 shRNA transduction was confirmed with fluorescent tags (GFP for G3BP2 and mCherry for G3BP1). SG KD was confirmed with immunostaining targeting G3BP1 and G3BP2 and imaged with confocal microscopy. AAV6 shRNA targeting both G3BP1 and G3BP2 was required for optimal SG KD when compared to G3BP2 shRNA alone, in which residual G3BP1 expression drove SG formation (**Supplemental Figure 3**).

*<u>Stress Granule Upregulation:</u>* Previous studies have shown that G3BP2 has a greater impact during SG formation than G3BP1.^85^ Therefore, we focused on triggering SG formation via G3BP2 upregulation. SG upregulation was achieved with a plasmid vector containing G3BP2 (pRP[Exp]-EGFP-CAG>hG3BP2, NM_001400006.1) from Vectorbuilder. iPSC-CMs were transfected with the G3BP2 plasmid via magnetofection in RPMI-C medium for 3, 6, 9, 24, and 48 hrs.^86^ Magnetofection technology works by attaching magnetic beads to the G3BP2 plasmid complexes; a magnetic field was then applied to concentrate the bead-plasmid complexes at the cell surface, facilitating cellular uptake. Plasmid transfection was confirmed with fluorescent tags (GFP). SG upregulation was confirmed with immunostaining targeting G3BP1 and G3BP2 and imaged with confocal microscopy. SG formation was quantified by SG number and average SG area (µm²). Upregulation of SG core protein G3BP2 alone was sufficient to trigger SG formation in iPSC-CMs (**Supplemental Figure 2**).

### Fluorescent Immunostaining

iPSC-CMs were replated on 35 mm glass-bottom dishes (#1.5) in preparation for staining. Samples were fixed with 2% paraformaldehyde (PFA) for 5 min at room temperature (RT), followed by a PBS wash (3 × 10 min at RT). Next, samples were permeabilized with 0.2% Triton X-100 in PBS (15 min at RT) and incubated with blocking buffer (1% bovine serum albumin, 0.1% Triton X-100 in PBS) for 2 hours at RT. Samples were labeled with primary antibodies (overnight at 4°C) and washed with PBS (3 × 5 min at RT). Following primary antibody labeling, samples were labeled with secondary antibodies (2 hrs at RT) and washed in PBS (3 × 5 min at RT). iPSC-CM dishes were maintained in PBS until imaging.

Proteins of interest were labeled with well-validated, commercially available primary antibodies: sarcomeric α-actinin [EA-53] (ACTN2) (1:1000, mouse monoclonal, Cat. #ab9465, Abcam), α-tubulin (1:500, rat µonoclonal, Cat. #sc-53029, Santa Cruz Biotechnology), G3BP Stress Granule Assembly Factor 2 (G3BP2) (1:500, rabbit polyclonal, Cat. #31799, Cell Signaling), G3BP2 CoraLite Plus 488 (1:200, rabbit polyclonal, Cat. #CL488-16276-100UL, Proteintech), G3BP Stress Granule Assembly Factor 1 (G3BP1) (1:2000, rabbit polyclonal, Cat. #13057-2-AP, Proteintech), L-type calcium channels (Cav1.2) (1:150, rabbit polyclonal, Cat. #ACC-003-200UL, Alomone Labs), and N-cadherin (N-cad) (4 µg/mL, sheep polyclonal, Cat. #AF6426, Bio-Techne). Samples were labeled with the following secondary antibodies: goat anti-mouse Alexa Fluor 405, goat anti-rat Alexa Fluor 568, goat anti-rabbit Alexa Fluor 647, goat anti-mouse Alexa Fluor 488, goat anti-rabbit Alexa Fluor 568, and goat anti-sheep Alexa Fluor 647 (1:4000; Thermo Fisher Scientific).

### Confocal Imaging

Confocal images of dissociated iPSC-CMs (**Figures 3, 5, and 6**), and isolated murine cardiomyocytes (**Figure 1, 2**) were acquired using a Leica SP8 LIGHTNING single-photon confocal microscope equipped with 5 solid-state lasers (405, 488, 514, 552, and 638 nm, 12 mW each), a 63×/1.4 NA oil immersion objective (120 nm resolution), 2 HyD GaAsP detectors, and 2 high-sensitivity photomultiplier tube detectors (Leica). Images were collected as z-stacks at Nyquist sampling rate or greater, as previously described^87^, and are presented as single optical planes. SGs were analyzed for number and average area (µm²) using the Analyze Particles function in ImageJ. Remaining structures were quantified as percent area in ImageJ. Colocalization between two structures was quantified using Manders’ colocalization coefficients with an ImageJ plugin (JACoP).^88^

### Optical Mapping

Whole-cell AP morphology was assessed by optical mapping. Low-density iPSC-CM cultures (control, arsenite (0.5 mM), G3BP2 upregulation, and SG knockdown; 60,000 cells/plate) were externally labeled with a far-red voltage-sensitive dye (BeRST-1) in Tyrode solution (in mM: 130 NaCl, 0.4 NaH₂PO₄, 5.8 NaHCO₃, 5.4 KCl, 0.5 MgCl₂·6H₂O, 1.5 CaCl₂·2H₂O, 25 HEPES, 22 glucose; pH adjusted to 7.3 with NaOH) and field paced at 1 Hz (10 ms bipolar pulses, 20 V) with graphite electrodes (MyoPacer, IonOptix, USA) at 37°C. Optical APs were recorded using a high-speed imaging system (Eclipse Ts2R, Nikon; Tokyo, Japan) equipped with a digital CMOS camera (Hamamatsu ORCA-Flash4.0 V2; Japan).^89^ Temperature was maintained at 37°C with a thermal plate integrated into the imaging stage (Tokai Hit; Shizuoka, Japan). Video data were analyzed with a custom open-source MATLAB script (available at: https://huebschlab.wustl.edu/resources-2/).^90^ Specifically, APs were characterized by the AP duration at 90% and 80% repolarization (APD90 and APD80). APD80 repolarization was used due to altered morphology during phase 4 of the AP.

### Statistical Analysis

Both structural and electrophysiological data were analyzed using GraphPad Prism software. Distribution was assessed using normality and lognormality tests. Differences in means were assessed by one-way ANOVA with multiple comparisons between groups (Welch’s correction was applied for non-normally distributed data or Kruskal-Wallis test was used for non-parametric comparisons). Differences in overall distribution for two-group comparisons were assessed by the Kolmogorov-Smirnov test. P values ≤ 0.05 were considered statistically significant. Varying degrees of significance are denoted in figures: *P ≤ 0.05; **P ≤ 0.01; ***P ≤ 0.001; ****P ≤ 0.0001.

### Study Approval

All animal procedures were approved by Institutional Animal Care and Use Committee at Washington University in St. Louis and performed in accordance with the Guide for the Care and Use of Laboratory Animals published by the U.S. National Institutes of Health (NIH Publication No. 85-605 23, revised 2011).

## Author Contributions

H.L.S, Y.D, R.P, and J.R.S conception or design of the work; H.L.S, A.Z.L, E.M, M.M.S, and A.K. performed research; N.H. contributed tools; C.C, S.G, R.M, J.M.N, K.J.L, D.M, D.M.O, and Y.D. contributed resources; H.L.S, I.F, T.M.S, and N.H. analyzed data; H.L.S, J.R.S, D.M.O, D.M, and A.K. edited the manuscript; H.L.S, and J.R.S. drafted the work or revised it critically for important intellectual content.

## Acknowledgements

The authors would like to thank Rebecca Mellor and Dr. Jeanne M. Nerbonne from the Department of Medicine and Developmental Biology for sample collection for this study, and Drs. Yifan Dai and Rohit Pappu for engaging discussions regarding biological condensates that helped shape the narrative of the work.

---

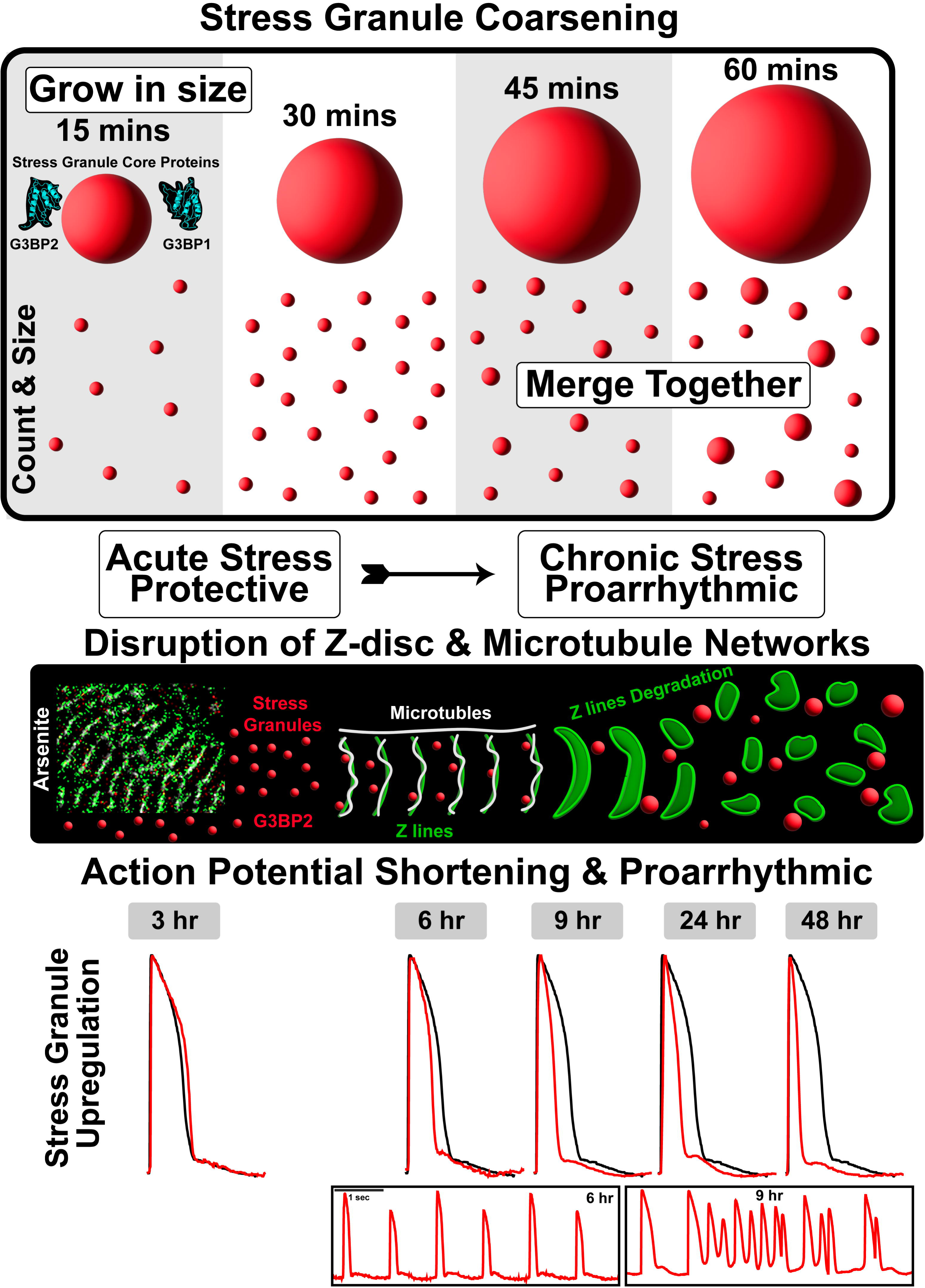

