## Supplemental Data for "Stress Granule Coarsening Is a Pathological Inflection Point for Cardiac Electrophysiological Dysfunction"

### Supplementary Figures

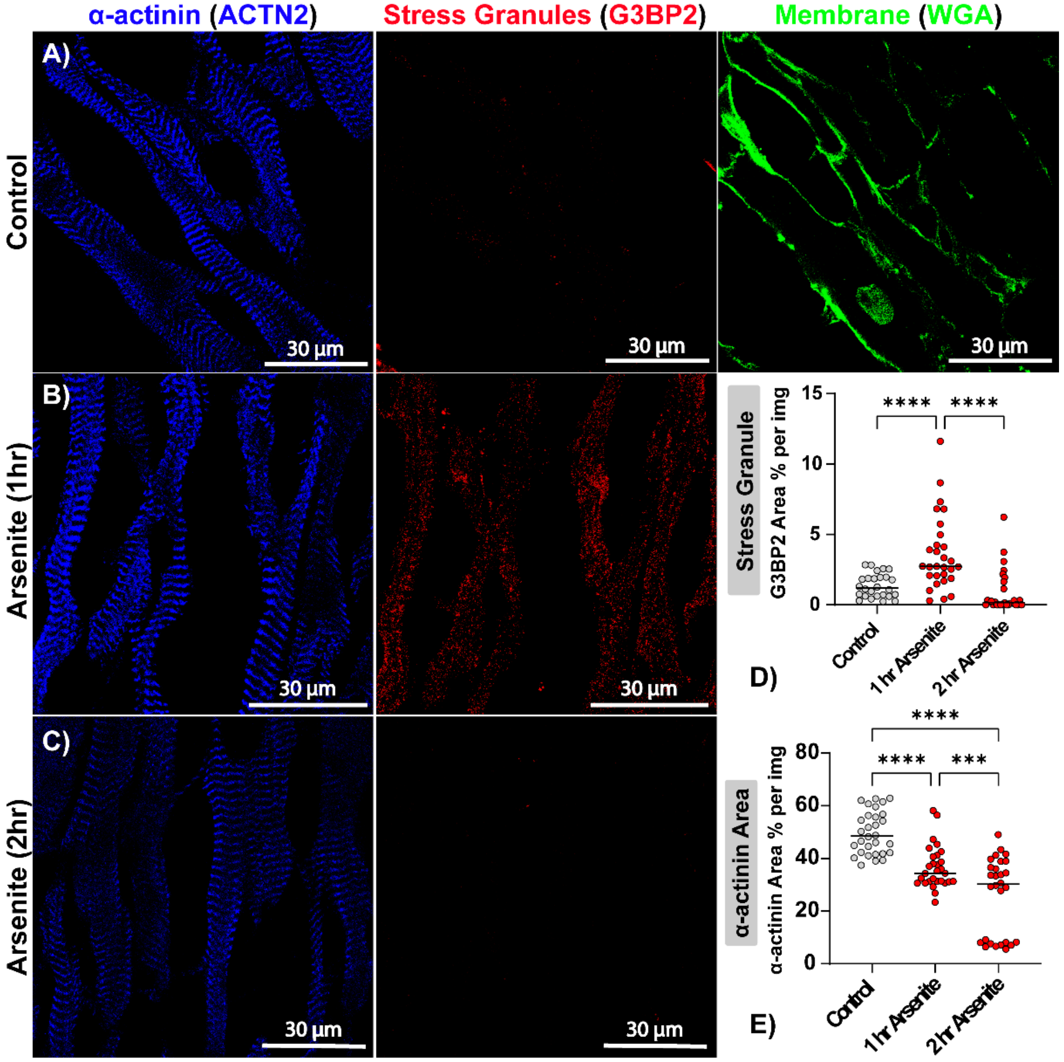


**Supplementary Figure 1. Acute Oxidative Stress in Whole Heart Murine Preparations.** Representative confocal images of cardiac tissue (5µm) slices from wild-type mice treated with acute oxidative stress (arsenite 0.5 mM) for 0, 1, and 2 hrs. **A-C)** Cardiac slices were stained for z-lines (α-actinin, blue) to track structural remodeling, cellular membranes (WGA, green) to track path of perfusion, as well as G3BP Stress Granule Assembly Factor 2 (G3BP2, red) for stress granule localization. SG formation was quantified by **D)** G3BP2 area per image (%). Structural remodeling was quantified as the **E)** percentage area of α-actinin. Murine studies consisted of 3 biological replicates with n ≥ 10 images per group. Differences in means were assessed by one-way ANOVA (NS = P > 0.05; *P ≤ 0.05; **P ≤ 0.01; ***P ≤ 0.001; ****P ≤ 0.0001).

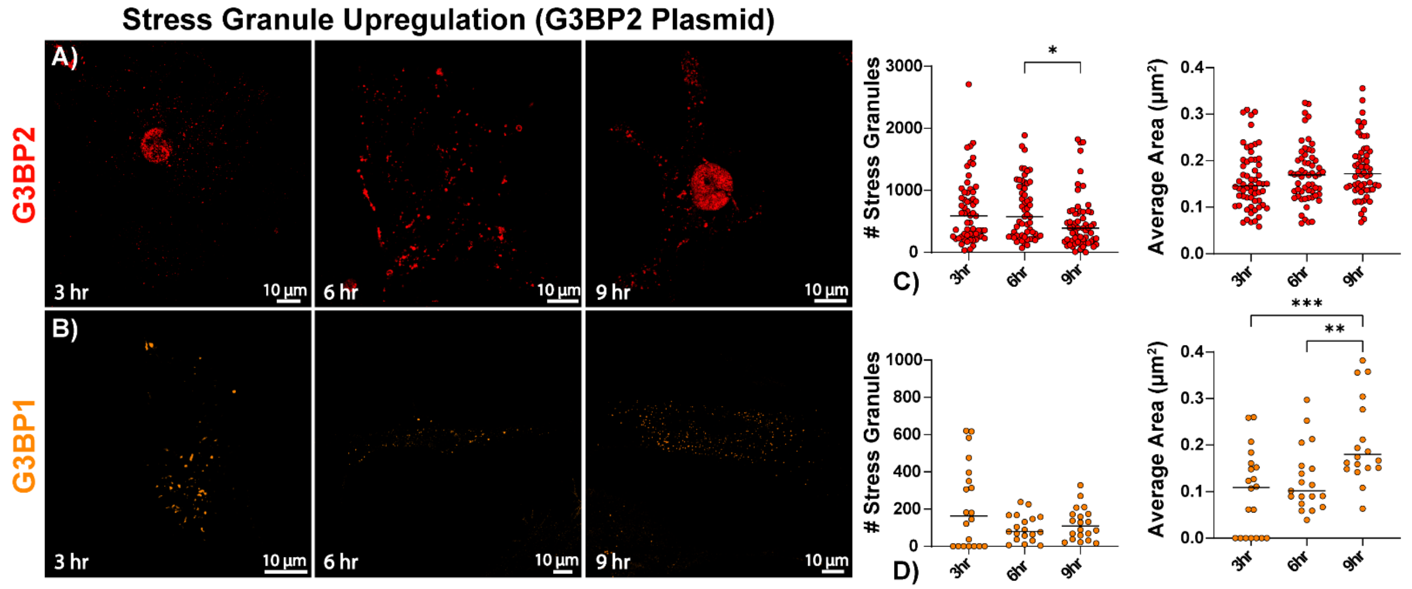


**Supplementary Figure 2. SG Upregulation in iPSC-CMs.** Representative confocal images iPSC-CMs treated with SG upregulation (G3BP2 plasmid) for 3, 6, and 9 hrs. **A-B)** iPSC-CMs were stained for G3BP Stress Granule Assembly Factor 2 (G3BP2, red, n = 60 cells) and G3BP Stress Granule Assembly Factor 1 (G3BP1, orange, n = 20 cells) for stress granule formation. SG formation was quantified by **C-D)** SG number and SG area (µm$^2$). Differences in means were assessed by one-way ANOVA (NS = P > 0.05; *P ≤ 0.05; **P ≤ 0.01; ***P ≤ 0.001; ****P ≤ 0.0001).

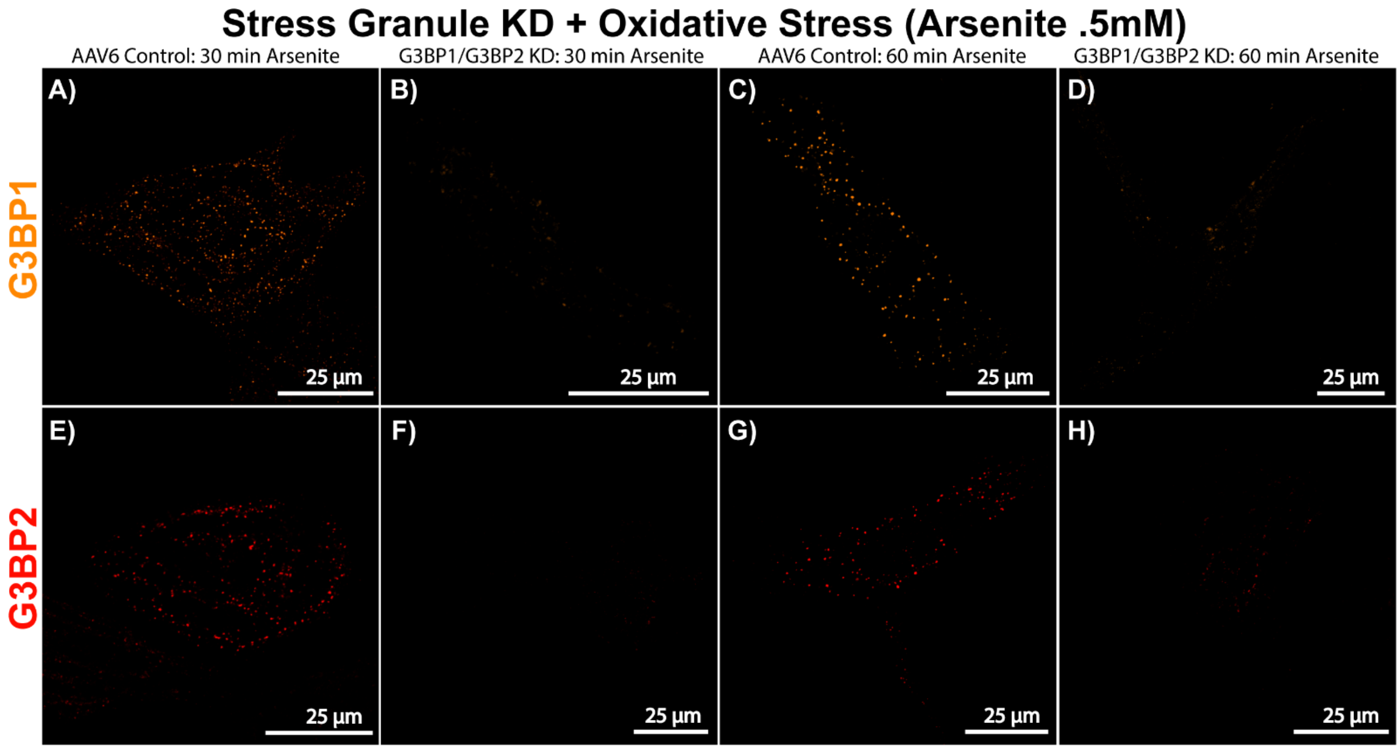


**Supplementary Figure 3. SG Knockdown in iPSC-CMs.** Representative confocal images iPSC-CMs treated with AAV6 control vector under **A, E)** 30 min and **C, G)** 60 min of oxidative stress and SG KD (G3BP1 and G3BP2) under **B, F)** 30 min and **D, H)** 60 min of oxidative stress. iPSC-CMs were stained for **A-D)** G3BP1 (orange) and **E-H)** G3BP2 (red) for stress granule formation.
